# The perception of odor pleasantness is shared across cultures

**DOI:** 10.1101/2021.03.01.433367

**Authors:** Artin Arshamian, Richard C. Gerkin, Nicole Kruspe, Ewelina Wnuk, Simeon Floyd, Carolyn O’Meara, Gabriela Garrido Rodriguez, Johan N. Lundström, Joel D. Mainland, Asifa Majid

## Abstract

Human sensory experience varies across the globe. Nonetheless, all humans share sensory systems with a common anatomical blueprint. In olfaction, it is unknown to what degree sensory perception, in particular the perception of odor pleasantness, is dictated by universal biological principles versus sculpted by culture. To address this issue, we asked 235 individuals from 9 diverse non-western cultures to rank the hedonic value of monomolecular odorants. We observed substantial global consistency, with molecular identity explaining 41% of the variance in individual pleasantness rankings, while culture explained only 6%. These rankings were predicted by the physicochemical properties of out-of-sample molecules and out-of-sample pleasantness ratings given by a separate group of industrialized western urbanites, indicating human olfactory perception is strongly constrained by universal principles.

## Introduction

In 1878 Margaret Wolfe Hungerford wrote that “Beauty is in the eye of the beholder”, suggesting that what one person finds beautiful, another may not. Similar to beauty, some argue that odor preference or valence is subjective—varying from person to person as well as across cultures (*1*–*7*). Fermented herring, for example, emits a smell described as the “most repulsive in the world,” but is a greatly appreciated delicacy in Sweden (*8*). At the same time, odor valence is considered to be the principal perceptual dimension by which odors are categorized and can be objectively predicted from chemical structure (*9*–*11*). It is unclear how to reconcile these perspectives: is odor preference entirely culturally relative or is it universally constrained by molecular structure? To address this, we designed a study to disentangle the influence of individual variability, cultural influence, and physicochemical properties on the perception of odor valence.

Previous studies attempting to address this question have been limited in a number of ways. Some studies take an ethnographic approach with traditional communities. These provide valuable insight into lesser-described cultures very different to western urbanites, but typically the studies are observational and focus on single odors that illustrate radical cross-cultural differences in odor preferences (*2, 6*). Other studies using an experimental approach have found universals in odor preferences; but these studies default to testing people with similar lifestyles and experiences—i.e., literate, educated, and technologically savvy individuals who partake of a common global fragrance and flavor industry (*12, 13*). In this study, a network of fieldworkers collected new experimental data regarding perceived odor pleasantness from nine diverse non-western communities sampled across varying subsistence styles, ecologies, and geographies. Critically, seven of these groups belonged to small-scale societies—including hunter-gatherers, horticulturalists, and subsistence agriculturalists—with a more traditional lifestyle and who do not experience the same chemical ecology as western urbanites (Fig. 1 and Methods).

**Figure 1.**
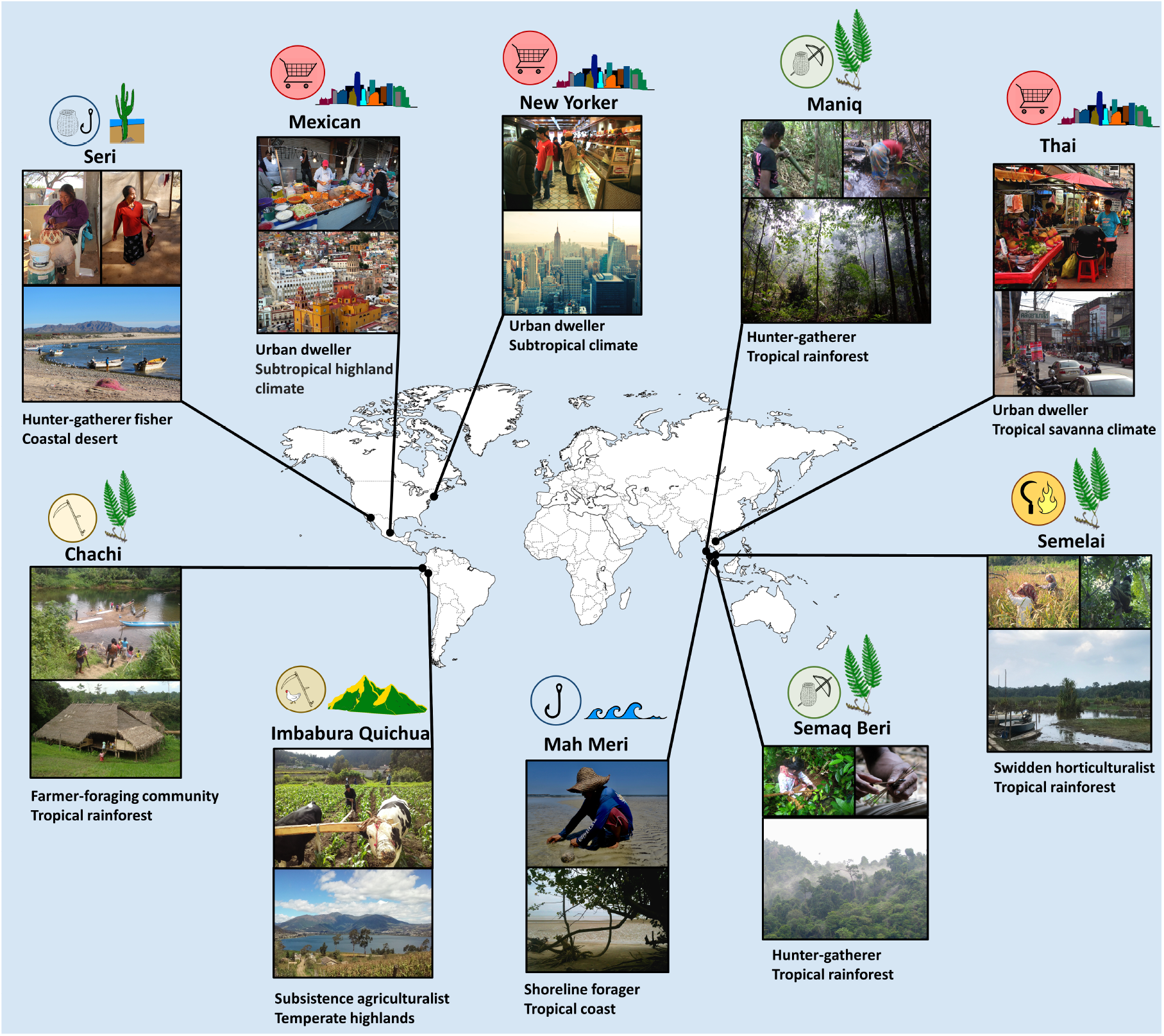
Cross-cultural sample. Odor preference ratings were taken from ten culturally and geographically diverse populations. These included the three hunter-gatherer groups, Seri, Maniq, and Semaq Beri, from a coastal desert and tropical rainforest respectively; one shoreline forager, Mah Meri, from a tropical coast; one swidden-horticulturalist, Semelai, from tropical rainforest; one farmer-foraging community, Chachi, from tropical rainforest; one subsistence agriculturalist, Imbabura Quichua, from temperate highlands; and three urban dwellers from industrial and post-industrial communities of bustling urban settings, Mexican, Thai, and American New Yorkers.

## Results

Odorants were selected based on a previous study with post-industrial urban dwellers from New York City (USA) who rated the pleasantness of 476 diverse molecules (*14*). We selected ten of these odorants such that the mean ratings would span the valence dimension from unpleasant to pleasant (for more details see Methods). Participants from the nine new communities were presented with ten pen-like odor-dispensing devices (*15*), each containing a unique odorant. A rank-order paradigm was chosen to assess odor pleasantness because not all groups had numeracy, and use of scales and ratings is not the norm in these communities. The pens were randomly ordered and placed in a line that faced the subject. The participant first smelled all the odors in front of them and then ordered the pens from most pleasant to most unpleasant (from their left-to-right).

If odor valence is universal, then all groups should rank odors in the same way. If, on the other hand, odor valence is learned from exposure to cultural traditions, then societies should differ in their perceived odor pleasantness, with a diverse set of rank orders across cultures. Using the within-culture mean ranking for each odorant, we found that odor valence rankings correlated strongly and positively across all cultures (Fig. 2, *r* = 0.82 ± 0.18), supporting the idea that culture has a relatively small influence overall on odor pleasantness.

**Figure 2.**
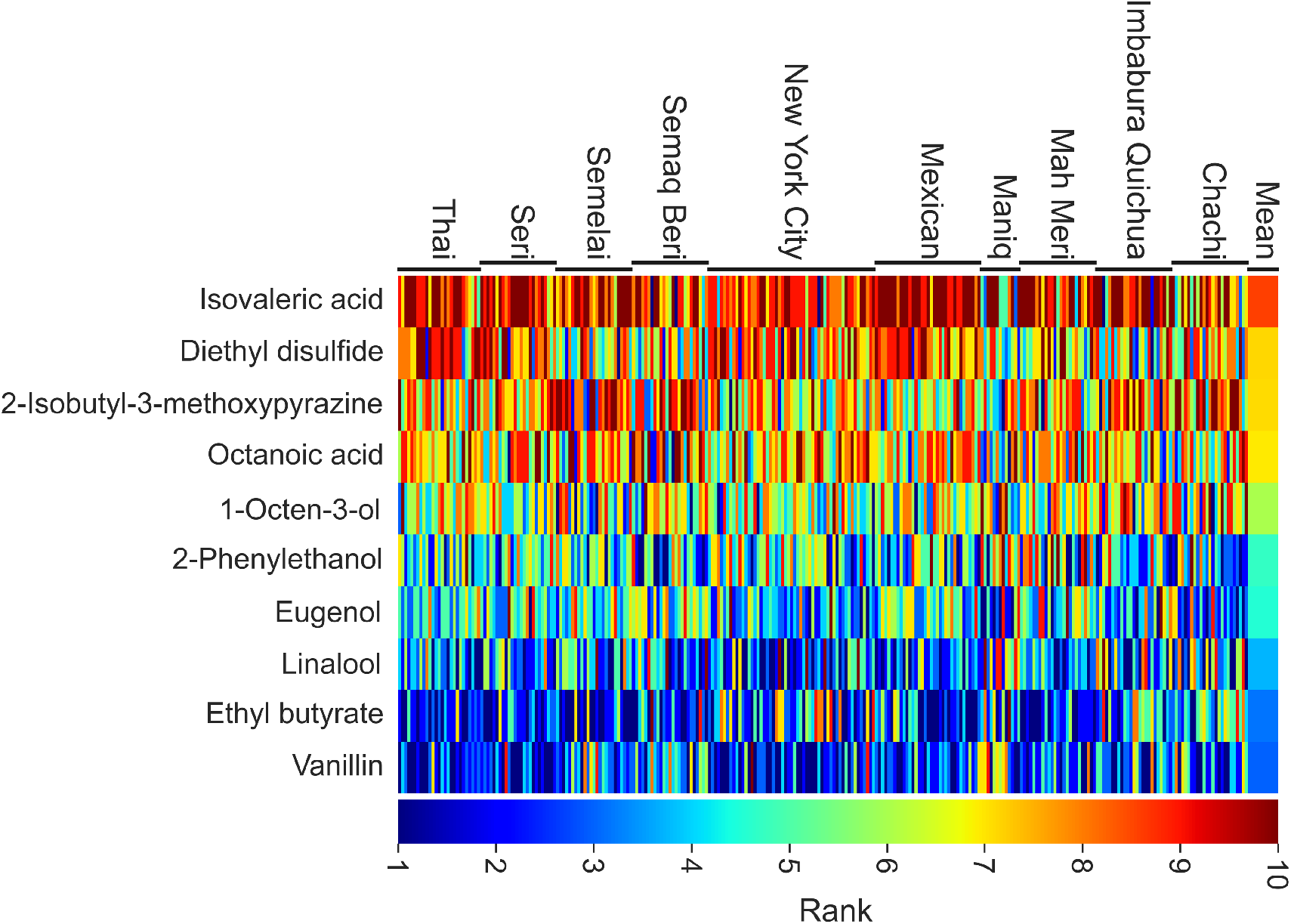
Pleasantness rankings across individuals and cultures. Between *n*=16 and *n*=55 individuals from each culture ranked each of 10 odorants in order from most (1, blue) to least (10, red) pleasant. Each color patch represents the integer ranking that one individual (from the culture indicated at the top) gave to one odorant (indicated on the left). The broad column on the far right represents the average ranking for each odorant across all individuals. Pleasantness rankings were also correlated for both the most pleasant and most unpleasant odorants (Supplementary Results Fig. S1).

We estimated the influence of culture, individual variability, and universal structure on odor valence using both frequentist and Bayesian statistical inference. The frequentist approach showed that a universal structure explained 41% of the variance in rankings, while culture explained only 6% (Fig. 3). The remaining 54% of the variance was due to individual variability, driven by some combination of individual preferences and perceptual noise. As a positive control, we simulated a case where culture drives odor preference by shuffling odor labels in a manner that was consistent for each member of a culture but varied across cultures. Under these conditions, 41% of the variance was explained by culture (Supplementary Results Fig. S2). This positive control shows that our method is sensitive enough to measure cultural variability should it exist. As a negative control for a possible effect of culture, we then shuffled individuals between cultures. Under these conditions, culture only explained 2% of the variance (Supplementary Results Fig. S2), not much smaller than the value observed in the unshuffled data. The analogous Bayesian model comparisons reached the same conclusions (Supplementary Results Fig. S3-S4). Consistent with only a small contribution for culture, direct assessment of inter-individual ranking similarity using Kendall’s tau showed that mean rank similarity for pairs of individuals within the same culture (tau = 0.32 ± 0.14) was only slightly higher than for pairs of individuals in different cultures (0.28 ± 0.11). In addition, a follow-up intensity ranking task showed that pleasantness ranking was not explained by the perceived intensity of the odorants (SI Appendix 1 and Supplementary Results Fig. S5). In summary, across the ten cultures we found only a weak contribution of culture to odor valence rankings.

**Figure 3.**
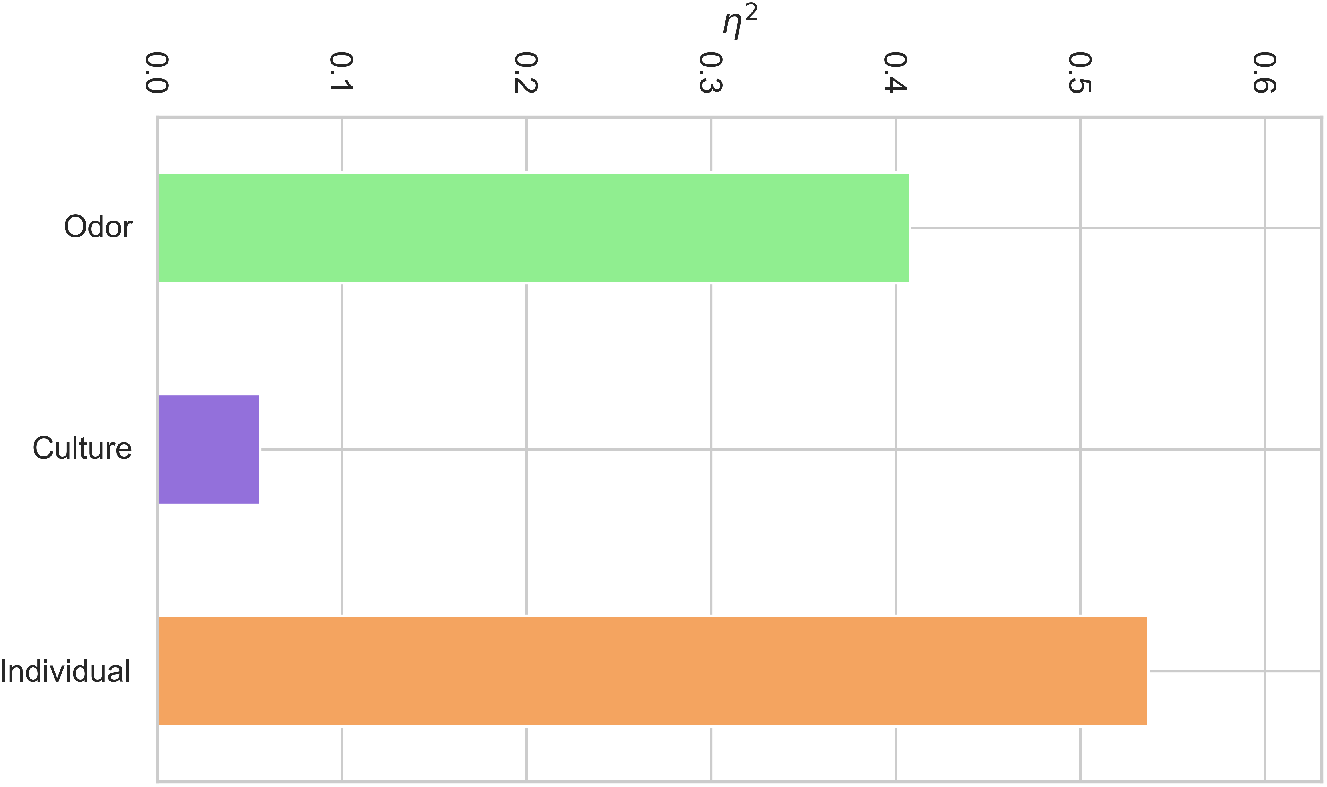
Effect size *η*^2^ from a two-way ANOVA for each of three factors that could potentially explain each individual’s pleasantness rankings. Culture membership (purple, 6%) plays a negligible role in explaining the variance in the observed odor pleasantness rankings; whereas Odorant identity (green, 41%) and Individual variability (orange, 54%) explain more.

If odor valence is mostly universal, then it should be possible to predict odor valence directly. Specifically, if physicochemical properties of odorants are the primary determinant of odor valence, the mean rank order from each culture should be predictable by a model trained using valence assessments made by a single culture. To do this we used the remaining 466 odorants from the original New York City data—excluding our 10 test odorants—to build a predictive model using the best random-forest algorithm from the DREAM Olfaction challenge applied to computed physicochemical features of each molecule (*16*). We then computed the rank order similarity between all pairs of individuals (including the model) using Kendall’s tau. For each and every culture, the within-culture mean rank order was more highly correlated with predictions from the model (on the test 10 odorants) than with any random participant from the same culture (Fig. 4). In other words, a universal model trained on responses of western urbanites to an independent set of odorants was at least as good a predictor of the culturally and ecologically diverse field data we collected here than data from the same culture and same set of odors.

**Figure 4.**
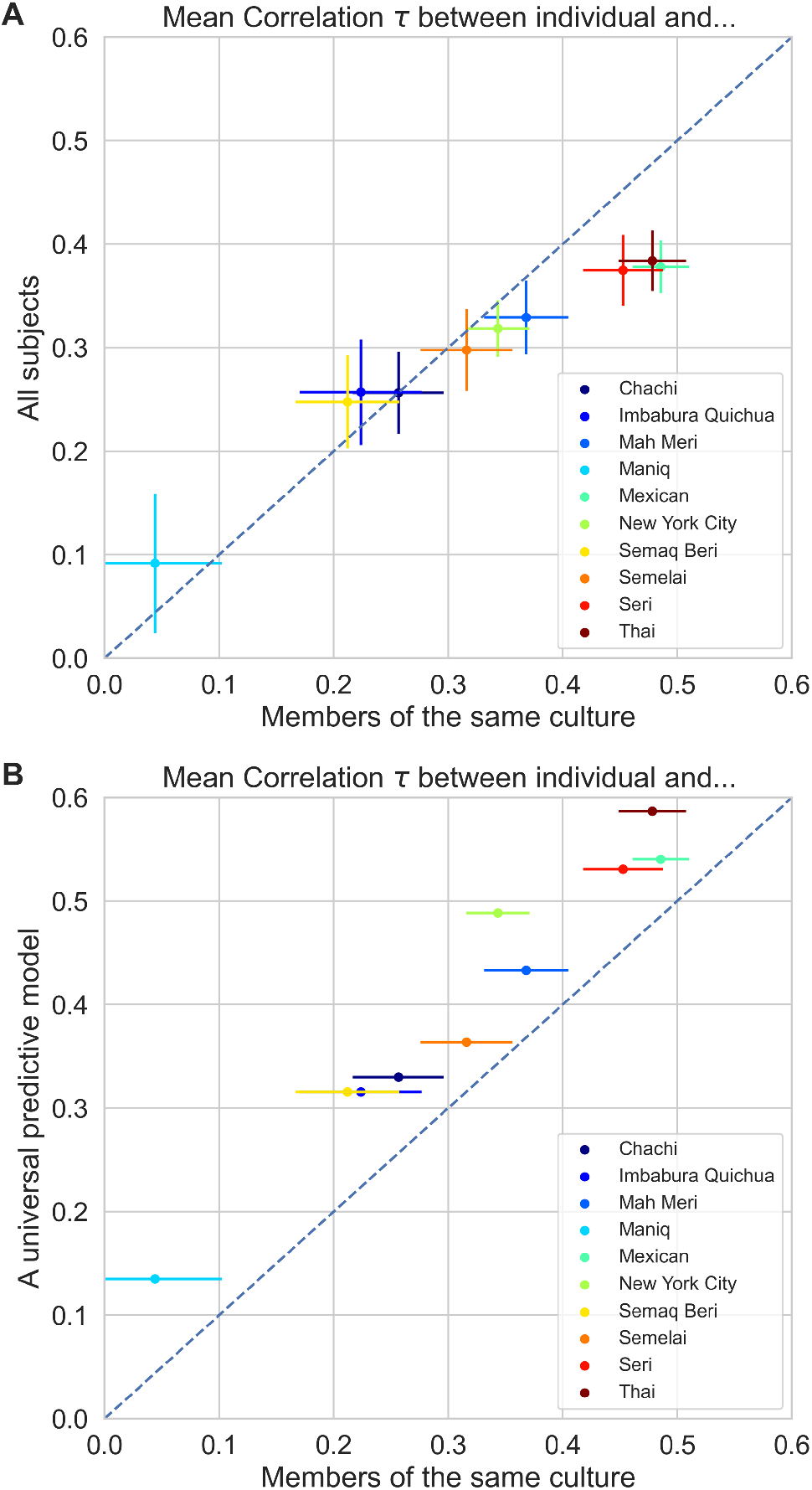
A universal model for odor pleasantness explains individual odor preferences. (A) The correlation of odor pleasantness rankings (Kendall’s τ) between each individual and other individuals from their culture (x-axis) is similar to the correlation between each individual and the entire population studied here. The hunter-gatherer Maniq showed the lowest correlation to other groups and to each other, with no cultural consensus. Critically, the Maniq do not demonstrate a systematic alternative cultural odor preference; merely high levels of individual variation. A control task showed this was not because they misunderstood the ranking task (Supplementary Results Fig. S6). Differences in individual agreement across cultures could be explained by differences in the reliability of the instrument at different locations. (B) Rankings predicted by a computational model trained on perceived pleasantness ratings for out-of-sample odorants are more correlated with individual rankings for the odorants used here than are other individuals from the same culture.

## Discussion

Taken together, our results demonstrate that odor valence perception is largely independent of cultural factors such as subsistence style and ecology, and can be predicted from physicochemical properties. This is striking, and contrary to what would have been predicted from a cultural relativity perspective (*1*–*7, 17, 18*). While it is widely accepted that valence is the principal perceptual axis of olfaction (*9*–*12, 19*), there has also been wide support for the idea that most aspects of olfactory perception are highly malleable and mainly learned (*1*–*7, 17, 18, 20, 21*), and importantly have little to do with an odorant’s physicochemical properties (*22*). Odor pleasantness is demonstrably plastic and modulated by factors such as early exposure (*23, 24*) and context (*25*–*30*). Our data do not adjudicate between learned versus innate explanations of odor pleasantness perception. Global regularities in odor perception could indicate common and shared experiences across all human groups. Infant data from diverse cultural contexts could adjudicate between these possibilities; although even here there are challenges since the fetus is already being enculturated into a specific chemical environment (*31*).

Nevertheless, our data demonstrate that physicochemical structure, rather than culture, seems to be the primary predictor of the pleasantness of most odors. This is also reflected in the fact that odor valence is shared across a wide range of species, possibly due to processes at the receptor level that may shape the valence for monomolecular odorants as well as complex mixtures (*32*–*36*). Critically, we show there is a universal bedrock of olfactory perception that is shared among all people.

## Materials and Methods

### Sample

Our sample consisted of data from 10 communities with diverse modes of subsistence living in varied environments (Fig. 1). The data were collected and treated according to the ethical guidelines of the American Psychological Association, and the protocol was approved by the Ethics Assessment Committee at Radboud University. Informed consent was obtained in writing or orally as appropriate to each community. We briefly describe each community in turn.

#### Semaq Beri

The hunter-gatherer Semaq Beri live in the northeast of the Malay Peninsula. Traditionally they moved about the tropical rainforests in small bands of eight to ten families, making temporary camps of lean-to shelters, hunting and fishing, and foraging for the many kinds of wild tubers and seasonal fruits. They have become increasingly sedentized since the establishment of resettlement villages in the mid-1970s. The participants in this study live in a village of around 300 people, and maintain a forest-based subsistence mode. They speak the Semaq Beri language which belongs to the Austroasiatic language family. The total Semaq Beri population is approximately 2,300. Sample: There were 25 subjects (13 female and 12 male, *M*_age_ = 33.3 years, *SD* = 14.3 years) a number that is equivalent to approximately 1% of the total Semaq Beri population.

#### Maniq

The Maniq inhabit a mountainous region in the interior of isthmian Thailand. The area is covered by tropical evergreen forest. Maniq subsistence is hunting, gathering, and exchange of forest products for food. The Maniq population is around 300 with the size of a residential group varying day-to-day, usually close to 25-35. The group lives in temporary camps in the rainforest with minimal material possessions. Maniq is the main and first language, although everyone can understand and speak Southern Thai (with varying degrees of proficiency). Only a handful (<5) of Maniq have received basic schooling and most are illiterate. Sample: There were 16 subjects (8 female and 8 male, *M*_age_ = 33.4 years, *SD* = 12.5 years) a number that is equivalent to approximately 5% of the total Maniq population.

#### Seri

The Seri are a traditionally hunter-gatherer-fisher, semi-nomadic people, but since the mid-20th century they are now more sedentary. They take part in small-scale fishing operations, small ecotourism enterprises, sell permits to hunt on their land, work as hunting guides and benefit from the sale of arts and crafts, especially baskets made of desert limberbush. Seri live in 2 small villages in northwestern Mexico, along the coast of the Gulf of California in the Sonoran Desert. Their traditional homeland includes the biggest island in Mexico, Tiburon Island. The population size is between 900 and 1,000. The Seri people speak the Seri language. The participants in this study were from the village El Desemboque de los Seris, Sonora. Sample: There were 25 subjects (19 female and 6 male, *M*_age_ = 39.3 years, *SD* = 16.4 years) a number that is equivalent to approximately 2.5% of the total Seri population.

#### Semelai

The Semelai reside in the southwest of the Malay Peninsula in an area of lowland tropical rainforest. Traditionally they dwelt in small family groups scattered throughout the forest, growing rice in swiddens, fishing and hunting, and collecting forest products like rattan and resin for trade. Today the Semelai live primarily in resettlement villages of a few hundred people, and most are small-holder rubber tappers. They continue to fish and hunt. The Semelai speak the Austroasiatic language Semelai. Their total population is around 5,000. Sample: There were 25 subjects (13 female and 12 male, *M*_age_ = 38.3 years, *SD* = 13.8 years) a number that is equivalent to approximately 0.5% of the total population.

#### Mah Meri

The Mah Meri reside on the southwest coast of the Malay Peninsula in a rural landscape that has been dominated by rubber and palm oil plantations since the early 1900s. Traditionally the Mah Meri engaged in shoreline foraging along the mangrove-lined coast, hunting in the forest, and growing rice and other subsistence crops around their homesteads. Resettlement villages of several hundred people were founded in the mid-20th century. Cash-cropping was introduced, first coffee then palm oil, but the scarcity of land has long caused people to seek external employment, while others fish, or forage the shoreline. They speak Mah Meri, an Austroasiatic language. There are around 3,500 Mah Meri people. Sample: There were 25 subjects (13 female and 12 male, *M*_age_ = 39.4 years, *SD* = 15.7 years) a number that is equivalent to approximately 0.7% of the total Mah Meri population.

#### Imbabura Quichua

Imbabura Quichua people live in agricultural communities, planting crops like corn and potatoes, but are also famous for their long historical tradition of weaving which has developed into an important handcraft industry. Like the other Highland Quichua people of Ecuador, they speak a local variety of the Ecuadorian Highland Quichua language descended from the Quechua language introduced by the Incas from modern Peru. While many Imbabura Quichua people maintain a traditional rural lifestyle, eat local food, and speak mainly Quichua, others are connected to the national and overseas economies through trade, travel, and the tourism industry, and are bilingual in Spanish; both types of participants were included in the study, conducted in a semi-rural, semi-urban area. Sample: There were 25 subjects (14 female and 11 male, *M*_age_ = 43 years, *SD* = 15.7 years) a number that is equivalent to approximately 0.0004% of the total Imbabura Quichua population of approximately 60,000.

#### Chachi

Traditionally the Chachi lived in isolated homesteads but today they live in small communities along the Cayapas river and its tributaries. Their lifestyle is mainly based on subsistence agriculture, with plantain as the basic staple, in addition to fishing and hunting. They also plant cash crops like cacao and engage in other activities like basketwork and logging, and use income to purchase outside supplies such as white rice, which has become an important staple in recent years. Their language is called Cha’palaa, from the Barbacoan language family. Participants are from a remote rural area where local people maintain a relatively traditional and autonomous lifestyle in which Cha’palaa is the dominant language and Spanish is used by a minority who have some experience outside the community. Sample: There were 25 subjects (13 female and 12 male, *M*_age_ = 44.6 years, *SD* = 14.9 years) a number that is equivalent to approximately 0.0025% of the total Chachi population of about 10,000.

#### Mexican

Mexico is a country with a population of 126 million. Mexico is considered to be ethnically diverse. Group residence varies considerably with large cities having many millions whereas small towns can have populations in the thousands or less. Our sample of subjects came from Mexico City which has a population of approximately 8.9 million people. The majority of the participants of the study were university employees (e.g., office workers). All subjects had access to all modern technologies (e.g., internet and television). The subjects were tested in Mexican Spanish. Sample: There were 35 subjects (19 female and 13 male, *M*_age_ =39.8 years, *SD* = 15.5 years) a number that is equivalent to approximately 0.000004% of the total Mexico City population.

#### Thai

Thailand has a population of almost 70 million with a mixture of ethnic groups. The data for this study was collected on the campus of the University of Ubon Ratchathani in Northeastern Thailand. The city of Ubon Ratchathani is a capital and an urban center of the province Ubon Ratchathani with 1.87 million inhabitants. Sample: The participants were from Ubon University and included university students and university employees (instructors, guards, cooks, shopkeepers). All subjects had access to all modern technologies (e.g., internet, television, etc.). The subjects were tested in Thai. There were 27 subjects (16 female and 11 males, *M*_age_ = 30 years, *SD* = 14.2 years) a number that is equivalent to approximately 0.000014% of the total of Ubon Ratchathani province.

#### New Yorkers

USA is a country with a population of 328 million. Group residence varies considerably with large cities having many millions whereas small towns can have populations in the thousands. Our sample of subjects came from New York City, a racially and ethnically diverse city, which has a population of approximately 8.4 million people. Sample: There were 55 subjects (33 female and 22 male, *M*_age_ = 34.6 years, *SD* = 9.5 years) a number that is equivalent to approximately 0.0000065% of the total New York City population.

### Materials

Stimuli were presented in Sniffin’ Sticks (Burghardt®, Wedel, Germany) permeated with the odor diluted in mineral oil. The ten odors, all from Sigma-Aldrich, used the DREAM Olfaction Prediction Challenge dilution series (1): Isovaleric acid (CAS 503-74-2; 1/100,000 volume/volume), Diethyl disulfide (CAS 110-81-6; 1/1000), Octanoic acid (CAS 124-07-2; 1/1000), 2-Isobutyl-3-methoxypyrazine (CAS 24683-00-9; 1/1000), 1-Octen-3-ol (CAS 3391-86-4; 1/1000), 2-Phenylethanol (CAS 60-12-8; 1/1000), Ethyl butyrate (CAS 105-54-4; 1/1000), Eugenol (CAS 97-53-0; 1/1000), Linalool (CAS 78-70-6; 1/1000), and Vanillin (CAS 121-33-5; 1/10).

### Procedure

Our main experimental protocol was a pleasantness rank order task. As a control, we also conducted an intensity rank order task. For the Maniq alone we conducted a protocol validation task using pictorial stimuli.

#### Pleasantness rank order task

The odors were randomly ordered on a holder so they made a line facing the subject. The subjects were told in their native language (i.e., Spanish, Seri, Imbabura Quichua, Cha’palaa, Thai, Maniq, Semelai, Semaq Beri, Mah Meri) to initially smell all the odors in front of them (briefly and one at a time), and after smelling all odors to order them from the most pleasant to the most unpleasant (their left-to right). The experimenter made sure that subjects did not smell an odor more than 2-3 seconds and that there was an interstimulus interval of 20 seconds between odor presentations. After subjects had smelled all odors on first encounter, they could freely re-sample the odors again while ranking them. To verify that subjects had understood the task correctly the experimenter asked them to point to the most pleasant and unpleasant odor after they finished the rank order task.

#### Intensity rank order task

Data collection was in two waves. In the first wave, data was collected from the Maniq, Seri, Mexican and Thai. We suspected some odors were vulnerable to the humid weather conditions in the Maniq site, one of the first to be tested, although we were not able to measure this objectively at the time. In a second wave of data collection, we collected data for judgements of odor intensity. The same participants from five of the nine groups (Chachi, Imbabura Quichua, Semelai, Semaq Beri, Mah Meri) that participated in the odor pleasantness rank order task also ranked odors by intensity using the same paradigm. The subjects were told in their native language that the task was first to smell all the odors in front of them (briefly and one at a time) and then to order them from the strongest to the weakest (their left-to right). As before, the experimenter made sure subjects did not smell the odor more than 2-3 seconds and that there was an interstimulus interval of 20 seconds between odor presentations. Before the start of the intensity rank ordering of the odors, the subjects undertook a control task to ensure they understood that they had to order odorants according to intensity and not odor pleasantness. In this control task, four different concentrations of the same odorant (i.e., 1-octen-3-ol in paraffin oil) were presented to the subject in steps of: 1/10,000,000 (basically blank); 1/100; 1/10; and 100% 1-octen-3-ol. To minimize adaptation effects, subjects first compared the two weakest concentrations, then the two strongest concentrations, then the second strongest and second weakest. Finally, they ordered odors from the strongest to the weakest (their left-to-right).

#### Ranking protocol validation for Maniq

As the Maniq are not accustomed to formal testing, we included a control to confirm that they could rank stimuli. We asked the same Maniq participants to rank order a set of 8 photographs of animals according to their hedonic value from most pleasant to most unpleasant (their left-to-right). We chose animals that varied in hedonic values based on long-term ethnographic fieldwork with the Maniq (Supplementary Results Fig. S6).

### Statistical Analysis

Full analysis details are provided in the Supplementary Analysis file. Code and data have been provided to the reviewers and will be made publicly available upon acceptance on GitHub at http://github.com/rgerkin/shared-pleasantness. Values reported in the main text are mean ± standard deviation. Correlations are reported using Pearson’s correlation (for means across groups) or Kendall’s tau (for comparisons between individuals).

## Acknowledgments

The fieldwork for this paper was funded by The Netherlands Organization for Scientific Research, NWO VICI grant “Human olfaction at the intersection of language, culture and biology” (Project number 277-70-011). This work was funded in part by the National Institutes of Health (U19NS112953, R01DC018455). A.A was funded by a grant from the Swedish Research Council (VR 2018-01603).

