## Supplemental Materials for "The perception of odor pleasantness is shared across cultures"

**This PDF file includes:**

Fig. S1-S7.

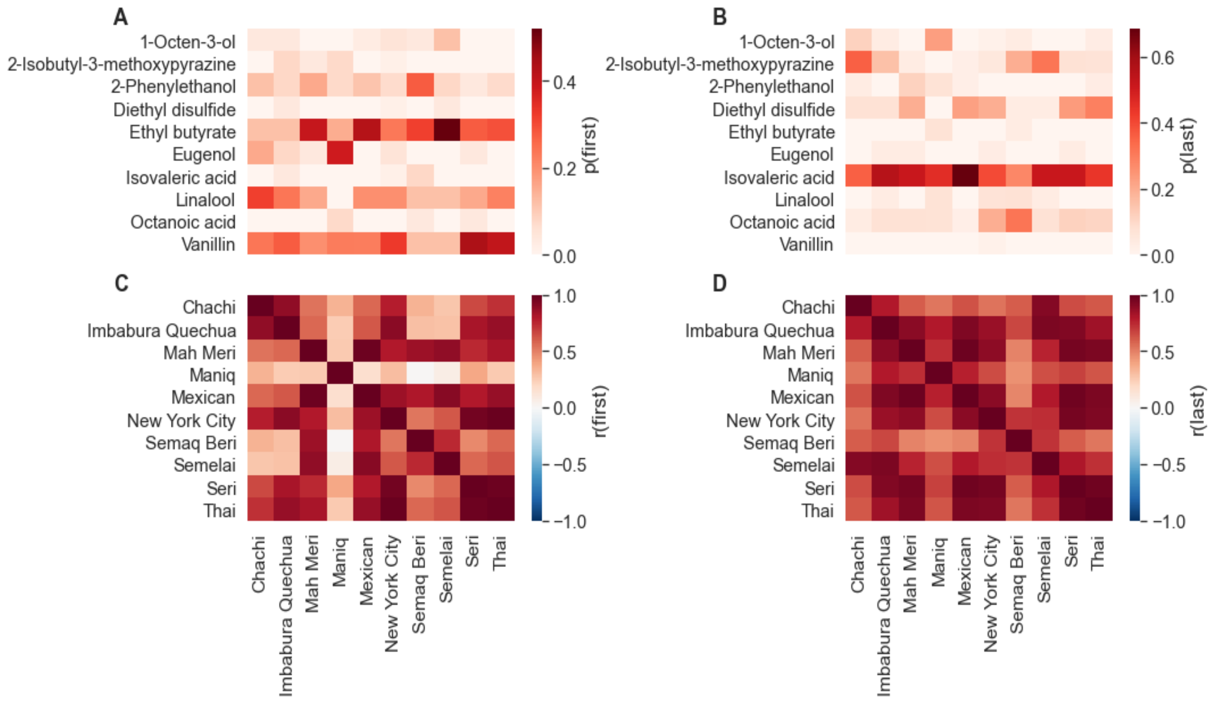


**Fig. S1. Comparison of highest-ranked and lowest-ranked odorants in each culture.** Pleasantness rankings are correlated for both the most pleasant and most unpleasant odorants. (A) Fraction of individuals within each culture that ranked the given odorant as the most pleasant of the 10. (B) Same as A, but for the least pleasant odorant. (C) Correlation matrix computed from panel A; this shows correlation between cultures in the number of individuals that rated each odorant as the most pleasant. (D) Same as C but showing a correlation matrix computed from B, for the least pleasant odorants.

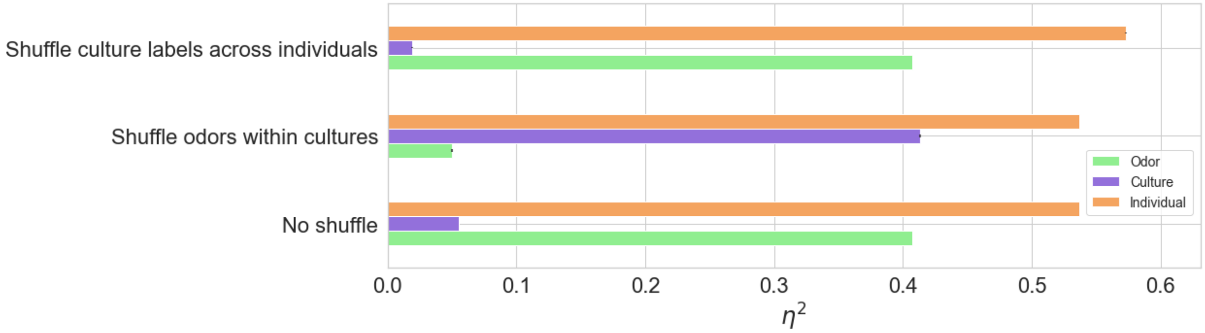


**Fig. S2. Positive and negative control analysis.** These simulations demonstrate that alternative scenarios would have been detected using this design. The bottom set of bars correspond to the actual data. The middle bars correspond to a culture-specific shuffling of odorant labels applied to each individual (positive control). The top bars correspond to a shuffling of culture labels across individuals (negative control).

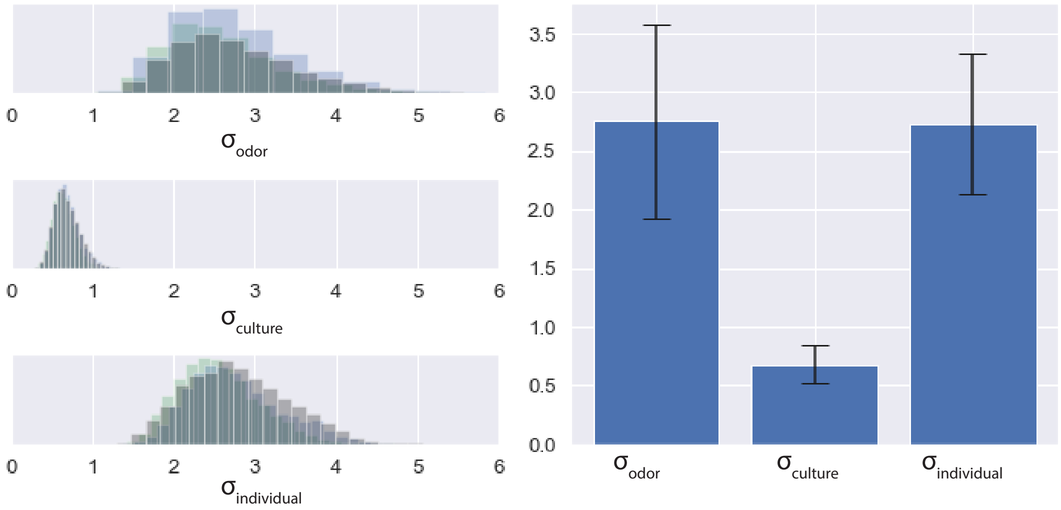


**Fig. S3. Bayesian model estimating the contribution of each factor to odor pleasantness.** A Plackett-Luce model (also known as exploded logit model or rank-ordered logit model) was constructed to perform full Bayesian inference on the rank data. In this hierarchical model, variance is partitioned between nested groups: individual, culture, and odor. The left-hand panel shows the marginal posterior distribution for the parameters associated with this variance. Each sampling chain (semi-independent estimate of the posterior distribution) is shown in a different color, but all estimates converge to roughly the same distribution. The right-hand panel shows the mean and standard deviation of these posterior distributions.

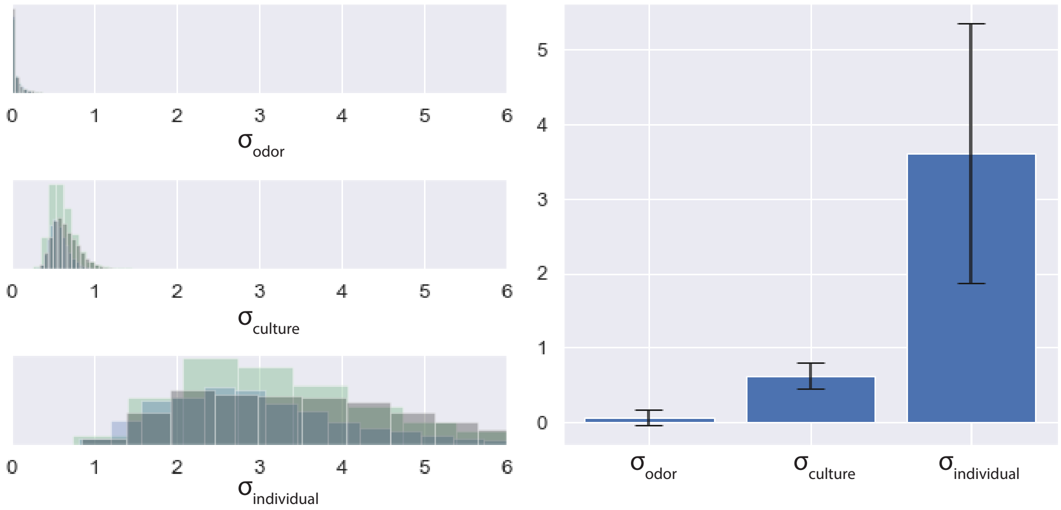


**Fig. S4. Bayesian positive control analysis.** Same as Fig. S3, but using a positive control in which we shuffled odor labels in a manner that was consistent for each member of a culture but varied across cultures. This destroys any universal structure to odor preference.

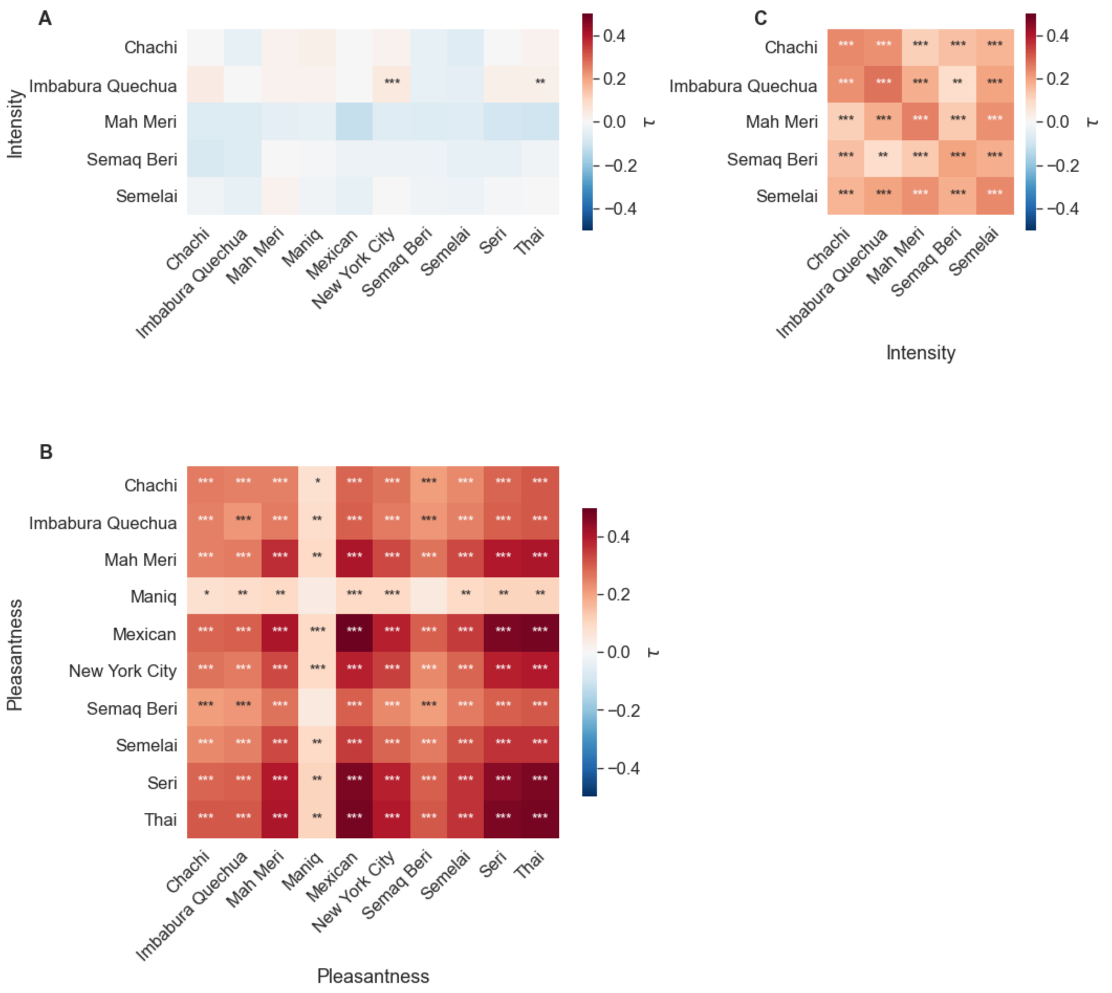


**Fig. S5: The correlations across cultures in pleasantness rankings are not explained by perceived intensity.** (A) Rank correlations (Kendall's τ) between individuals of the ranked pleasantness of the 10 odorants (x-axis) and the ranked intensity of the 10 odorants (y-axis) were computed, and the mean τ reported for such pairs of individuals within and across cultures. Such correlations were consistently near zero and not significant. Note that intensity data was collected in only some cultures, but from the same set of individuals as the pleasantness data. (B) The correlation between individuals for intensity alone was significantly positive, suggesting a "universal intensity" factor that is independent of pleasantness. (C) The correlation between individuals for pleasantness alone is positive and significantly greater than that observed for intensity. In all panels, comparisons between individuals and themselves were excluded. All rank correlations are annotated with statistical significance according to a binomial test, comparing the fraction of pairs of individuals that had $\tau>0$ against the null hypothesis of fraction ½, and applying a Bonferroni correction for multiple comparisons. *: $p<0.01$; **: $p<1e\text{-}4$; ***: $p<1e\text{-}10$.

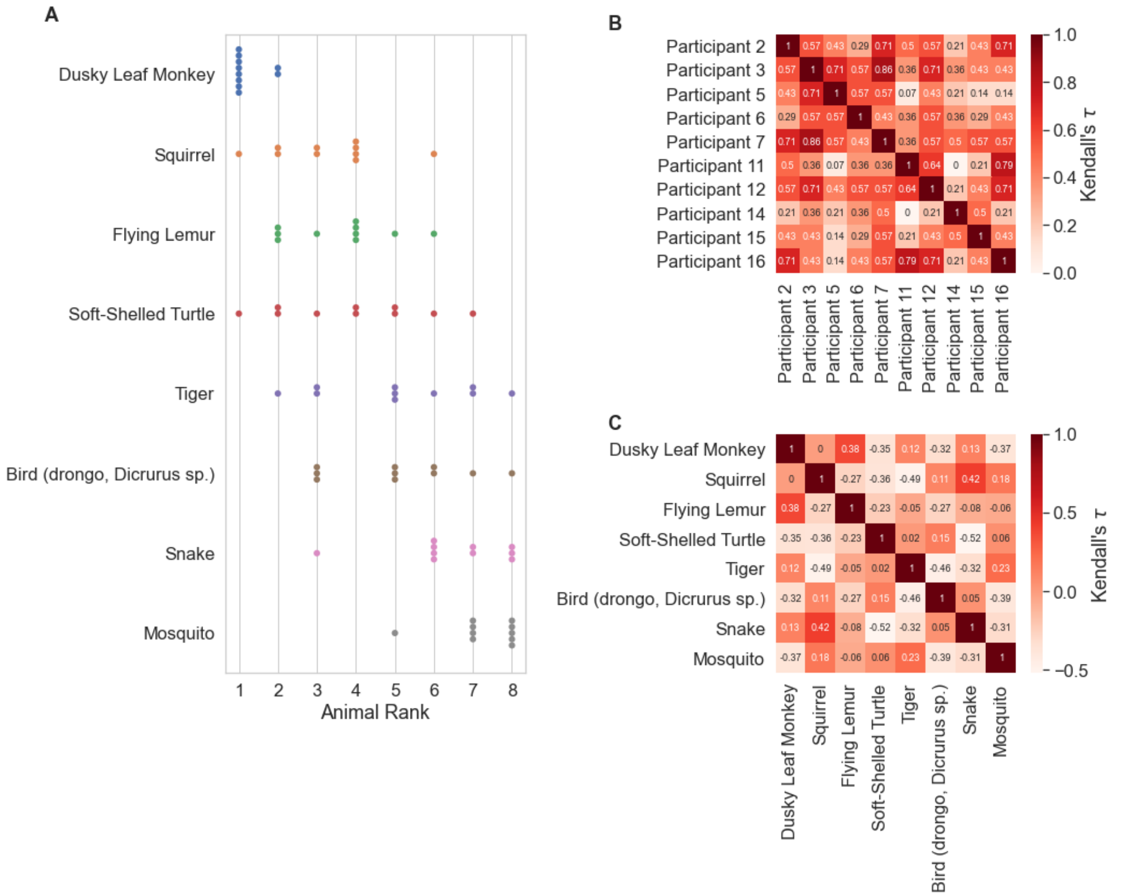

**Fig. S6. Maniq participants are capable of ranking items in an ordinal fashion.** Ten out of 16 Maniq participants were shown pictures of 8 animals and asked to rank them according to hedonic value (highest to lowest). (A) Each dot is the ranking given by one subject for one animal. (B) The rank correlation (Kendall's $\tau$ between participants). (C) The rank correlation between animals.
